# Formalin fixation and paraffin embedding interfere with preservation of optical metabolic assessments based on endogenous NAD(P)H and FAD two photon excited fluorescence

**DOI:** 10.1101/2023.06.16.545363

**Authors:** Adriana Sánchez-Hernández, Christopher M. Polleys, Irene Georgakoudi

## Abstract

Endogenous NAD(P)H and FAD two-photon excited fluorescence (TPEF) images provide functional metabolic information with high spatial resolution for a wide range of living specimens. Preservation of metabolic function optical metrics upon fixation would facilitate studies which assess the impact of metabolic changes in the context of numerous diseases. However, robust assessments of the impact of formalin fixation, paraffin embedding, and sectioning on the preservation of optical metabolic readouts are lacking. Here, we evaluate intensity and lifetime images at excitation/emission settings optimized for NAD(P)H and FAD TPEF detection from freshly excised murine oral epithelia and corresponding bulk and sectioned fixed tissues. We find that fixation impacts the overall intensity as well as the intensity fluctuations of the images acquired. Accordingly, the depth-dependent variations of the optical redox ratio (defined as FAD/(NAD(P)H + FAD)) across squamous epithelia are not preserved following fixation. This is consistent with significant changes in the 755 nm excited spectra, which reveal broadening upon fixation and additional distortions upon paraffin embedding and sectioning. Analysis of fluorescence lifetime images acquired for excitation/emission settings optimized for NAD(P)H TPEF detection indicate that fixation alters the long lifetime of the observed fluorescence and the long lifetime intensity fraction. These parameters as well as the short TPEF lifetime are significantly modified upon embedding and sectioning. Thus, our studies highlight that the autofluorescence products formed during formalin fixation, paraffin embedding and sectioning overlap highly with NAD(P)H and FAD emission and limit the potential to utilize such tissues to assess metabolic activity.

## 1. Introduction

Metabolic dysfunction is associated with numerous conditions such as cancer, diabetes, aging, neurodegenerative disorders, and cardiovascular disease [1-5]. Two-photon excited fluorescence (TPEF) can provide important insights regarding cellular metabolic function and dysfunction [6-10]. NAD(P)H (reduced nicotinamide adenine dinucleotide (phosphate)) and FAD (flavin adenine dinucleotide) are two naturally fluorescent co-enzymes involved in several metabolic pathways, such as glycolysis, oxidative phosphorylation, fatty acid synthesis, and oxidation and have been used to determine the oxido-reductive state of cells through the use of their TPEF intensity [8, 9, 11-13]. Since the fluorescence characteristics of NADH and NADPH are very similar, we use NAD(P)H to refer to potential contributions from NADH and/or NADPH [10]. Flavin contributions include fluorescence from free FAD, as well as FAD bound to LipDH, which is the most dominant contributor to the fluorescence of several cell types, and FAD bound to Complex II of the electron transport chain [10]. Flavin mononucleotide (FMN) bound to complex I may also contribute to cellular flavin associated autofluorescence. We will use FAD to refer to all of these flavin contributions. Additionally, the fluorescence lifetime of NAD(P)H can be used as another indicator of metabolic function as its bound and free states have distinct values [10, 14-17].

Given the potential of optical metabolic imaging, efforts have been largely focused on live cell or tissue imaging, both *in vivo* and *ex vivo* in freshly excised specimens [11, 18-25]. However, *in vivo* imaging is challenging and excised tissues cannot be preserved for large periods. Fixation is typically used to preserve the morphological structure and architecture of cells and tissues for an extended period [26]. Formaldehyde-based fixatives are the gold standard for histology and immunohistochemistry, and they are routinely used for histopathological assessment of tissue biopsies for medical diagnosis. These fixatives form cross links within and between proteins and nucleic acids [27]. Formaldehyde reacts with the free amino groups with a reactive hydrogen present in nucleotides and proteins, forming a reactive hydroxymethyl compound. This compound reacts with another reactive hydrogen from another amino group to form a methylene bridge cross-link [28]. Such cross-links may change the protein conformation, which in turn could impact their spectroscopic properties. Fixation is a progressive process that depends on time and temperature. Standard protocols establish minimums of 24-48 hours of fixation time for a 5 mm biopsy. Maximum times of fixation are usually not established unless antigen retrieval for immunohistochemistry is a concern. Formalin (10% Neutral Buffered Formalin, or NBF) is the most common formaldehyde-based fixative. Once the specimens are fixed, they undergo tissue processing (dehydration for paraffin embedding), sectioning, and staining protocols. Formalin fixed paraffin embedded (FFPE) tissues can be preserved for extended periods. Crosslinks formed during formalin fixation are a common source of tissue autofluorescence and methods have been developed to reduce its contribution in images [29, 30].

There are vast repositories of fixed tissue blocks, providing access to human tissues from patients with a number of conditions and following a wide range of treatments. However, the effect of the chemical and physical modifications of tissue due to fixation and downstream processing on the autofluorescence signals used for optical metabolic imaging is not fully understood and/or validated. Some studies have suggested that the metabolic interpretations from fresh tissue imaging can be maintained in tissue samples after fixation, both in the context of the optical redox ratio [31] and fluorescence lifetime [32, 33]. These studies have reported processing-associated alterations in the signals, both in terms of intensity and lifetime, as well as a spectral shifts. Nevertheless, these altered signals appeared to be correlated with the endogenous fluorescence of NAD(P)H and flavoproteins detected in the living specimens. Since some of these studies relied on cultured cells or employed mild fixation protocols and typically considered either intensity or lifetime based readouts, we sought to evaluate the impact of fixation, paraffin embedding and sectioning on both intensity and lifetime endogenous TPEF-extracted tissue metabolic function indicators. For this purpose, we acquired intensity and lifetime TPEF images at excitation/emission settings typically optimized for acquisition of NAD(P)H and FAD images from unstained squamous epithelial tissues that were freshly excised and compared them to images acquired from the same tissues after formalin fixation as well as after paraffin embedding and sectioning. We report on the impact of these processing steps on intensity fluctuations within a field, as well as on the ability to recover optical metabolic function readouts, such as the optical redox ratio, and the NAD(P)H short and long lifetime and bound fraction. The epithelial tissues are a suitable model for these studies as they include in the same specimen cell layers with subtle but reproducible metabolic function differences and we can monitor the ability to reproduce these differences during the different processing steps.

## 2. Methods

### 2.1 Samples

Murine cheek epithelial tissue biopsies were selected for imaging due to the anticipated variations in metabolic function across the depth of the squamous epithelia, which can serve to assess the maintenance (or loss) of the optical cellular metabolic profiles throughout the process of fixation and sectioning. The cheeks of three different rats were excised immediately after euthanasia. Each cheek was visually inspected and a piece of epithelium of approximately 5 by 5 mm was cut for imaging shortly after excision. Three ∼500 μm ink markings on the edges of the imaging surface (Fig. 1C) were placed in each biopsy for future identification of the region to be imaged after fixation and as a general reference for sectioning of the slides to ensure acquisition of histopathological sections of imaged regions from the bulk tissues. Fresh pieces were kept hydrated and placed in a glass bottom dish for imaging after marking. Once imaged, each biopsy was placed in formalin (10% Neutral Buffered Formalin) and imaged one week later. Samples were then dehydrated in successive concentrations of ethanol and xylenes, to replace the water with paraffin, resulting in FFPE tissue blocks as is established by standard protocols [34]. Sections (20 μm thick) contained within the three ink markings were mounted onto a slide, paraffin was removed with xylene and the sample was coverslipped. Images were acquired of the unstained FFPE sections (Figure 1).

**Fig. 1.**
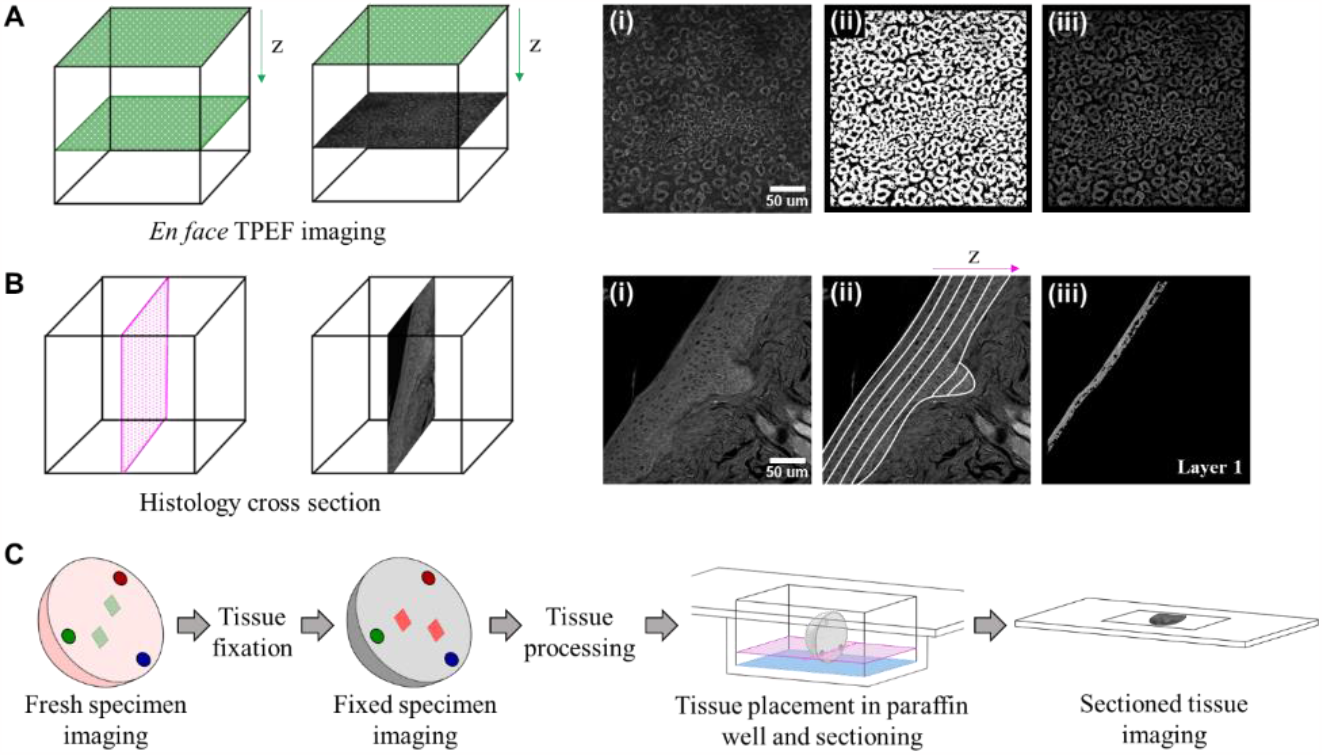
(A) Schematic of *en face* TPEF imaging of bulk tissues, with a TPEF sample image (i), mask used for isolation of cytoplasmic regions (ii), and resulting image for analysis (iii), respectively. (B) Schematic of histology cross section relative to *en face* TPEF imaging, with TPEF sample image of a cross section (i), division of the epithelium in order to analyze sections through depth (ii), and example of the superficial layer for analysis (iii). (C) Imaging workflow with tissue markings (circles) for acquisition of ROIs (squares) contained in the same regions of the tissue in freshly excised specimens, bulk fixed specimens, and sectioned specimens.

### 2.2 Imaging

Imaging was performed with a Leica TCS SP8 system equipped with a fs tunable laser (Insight, Spectra-Physics). All images were acquired with a 40X water immersion objective with a numerical aperture of 1.1. Fluorescence lifetime data was acquired with a commercial time correlated single photon counting module (TCSPC, PicoQuant). Endogenous fluorescence intensity of NAD(P)H was acquired with an excitation wavelength of 755 nm and a detection band of 460/50 nm (I_755/460_). FAD images were acquired at 860 nm excitation with a 525/50 nm detection band (I_860/525_). The redox ratio (FAD/(NAD(P)H+FAD)) assessments from images acquired using these studies have been validated with mass spectrometry experiments for squamous epithelial tissues [35]. We will use the term optical ratio to refer to the corresponding excitation/emission settings for these images when referring to processed tissues, i.e. (I_860/525_/(I_755/460_ + I_860/525_). Intensity image acquisition was made with 8 frame accumulation of photon counts at a 600 Hz rate for each frame line with a pixel dwell time of 0.4 μs. For the 860 nm excitation wavelength, second harmonic generation images were acquired with a 430/24 nm detection bandpass filter. This was used for masking of non-cellular regions in all images. For intensity analysis, three to four image stacks of 60-100 μm depth with a 4 μm step size were taken from each sample for each wavelength. For FLIM acquisition of NAD(P)H at the same emission and excitation bands, 4-5 evenly spaced depths were imaged along the entire thickness of the epithelium in each of the image stacks; data was acquired for one minute at each depth. For the FFPE unstained sectioned samples, three to four ROIs were imaged from each of the three samples. Figure 1C presents a schematic of the imaging workflow of each tissue sample in its different conditions. Tissue markings with a needle, depicted with circles in the schematic, were made along a region of epithelium in the freshly excised sample. All ROIs imaged, represented as squares, were contained within the markings. Tissue was placed in 10% NBF for fixation and imaged later with ROIs contained again within the markings. Tissue was processed with standard protocols and placed in the paraffin well, such that two of the ink markings were contained in the same sectioning plane (shown as a blue surface in the schematic), thus, ensuring that the acquired histology sections (pink cross-section) belonged to the same region contained by the ink markings. Spectral images of an optical section with a full layer of cells were acquired at 755 nm and 860 nm excitation using the de-scanned detectors of the microscope with constant gain and power between emission wavelengths. Detection was performed from 400 to 700 nm for I_755_ and from 400 to 750 nm for I_860_ with a bandpass of 10 nm for each step in both cases. Intensity at each step was integrated and spectra were normalized to each sample’s maximum intensity.

### 2.3 Data processing and analysis

Images were normalized to account for different laser powers as described previously with the quadratic relationship between power and intensity [36], PMT gain was kept constant consistent with the photon counting mode. Downsampling of the images (1024 x 1024 pixels to 512 x 512 pixels) was performed to increase the distribution density of redox values while maintaining a pixel size resolution of 0.568 μm. A combination of filters (two Gaussian bandpass, and one Butterworth filter) was used to select the cell cytoplasmic regions and exclude the nucleus and background as previously established by our group [22]. Images were masked to remove regions with collagen signal to remove any contributions from non-cellular regions. Coregistration of the I_860/525_ images relative to the I_755/460_ images was done with the normxcorr2 function in MATLAB (R2019b v9.7.0) and a pixel-wise redox ratio map was created for each optical section calculated using the redox ratio definition of FAD/(FAD + NAD(P)H), as originally established by Chance and his colleagues [37]. For the depth-dependent analysis of the sectioned samples, 5-7 cell layers approximately 10 μm thick were identified within each epithelium and analyzed using the same methods as the optical sections of the bulk tissue images. These layers were created since the sectioned samples were cross sections as opposed to the *en face* images of the bulk epithelial tissues (Figure 1(A-B)).

To analyze the intensity variations of the I_755/460_ images, the power spectral density (PSD) was calculated as the squared amplitude of the 2D Fourier transform [38]. To account for the prevalence of relevant cellular and intracellular feature sizes, the area under the PSD curve was calculated for spatial frequencies between 7 x 10^−3^ and 2 x 10^−1^ 1/pixel. This analysis was only done with the bulk freshly excised and fixed tissue images, but not with the sectioned tissue, since the cross sections contained all layers of the epithelium in a single frame and imaged from a different perspective; thus, the comparison was not meaningful.

### 2.4 Fluorescence lifetime imaging analysis

Analysis of fluorescence lifetime images was done via the phasor approach which provides a graphical representation of each pixel in the lifetime image in phasor space [39]. Single mono-exponential decays were mapped along the universal semi-circle, while spectra with more than one lifetime were mapped inside the circle (see phasors in Figures 6 and 7) [39, 40]. Mapping all pixels of the image onto phasor space yielded an ellipse. An umbelliferone solution was imaged for calibration purposes. A bi-exponential decay was assumed for NAD(P)H, with pixels containing contributions from two different states of NAD(P)H: bound and free, associated with a long and short lifetime, respectively [41-43]. A line fit to the major axis of the ellipse crossed the universal semi-circle at locations that corresponded to the long and short lifetime values of NAD(P)H. For a given phasor located at the centroid of the 2D histogram, the distance to the long lifetime estimate divided by the total length of the fitted line represented the fractional intensity contribution of the long lifetime (long lifetime intensity fraction or LLIF), or the NAD(P)H bound fraction when NAD(P)H was the source of signal. This was calculated for every phasor. For phasors not located along the line, the LLIF was estimated by projecting each phasor to the fitted line.

The same filters as in the intensity image analysis were used to isolate the cytoplasmic regions and to eliminate contributions from the nuclei and background using the integrated intensity from the FLIM NAD(P)H acquisition. Signals associated with collagen were masked using the second harmonic generation images. For the sectioned samples analysis, masks were created in the same way as for the intensity depth-dependent analysis, but analyzed to extract TPEF lifetime associated metrics.

### 2.5 Statistics

Statistical analysis was performed with JMP Pro 16. A mixed-model ANOVA with post hoc Tukey tests for multiple comparisons was used to compare data with more than two groups (FLIM analysis). A Student t-test was used for the area under the PSD curve. Significance was established at p < 0.05.

## 3. Results

### 3.1 Tissue fixation and downstream processing yield autofluorescence that masks intensity-based optical metabolic readouts

Representative NAD(P)H (I_755/460_), FAD (I_860/525_), and corresponding redox ratio coded images for the fresh tissues are shown in Fig. 2. Images acquired for the same excitation and emission conditions are shown from fixed and sectioned tissues. Higher levels of I_755/460_ signal overall and a lower contribution to the I_860/525_ channel are observed in the fresh tissue specimens. Fixed specimens, especially those sectioned, have a higher contribution on the I_860/525_ channel and a lower intensity in the I_755/460_ channel. These signal levels are reflected in the optical ratio (I_860/525_/(I_755/460_ + I_860/525_) coded images (figure 2(G-I)) with bluer hues representing lower values for fresh tissue compared to the yellow/red hues of the fixed bulk and sectioned tissue.

**Fig. 2.**
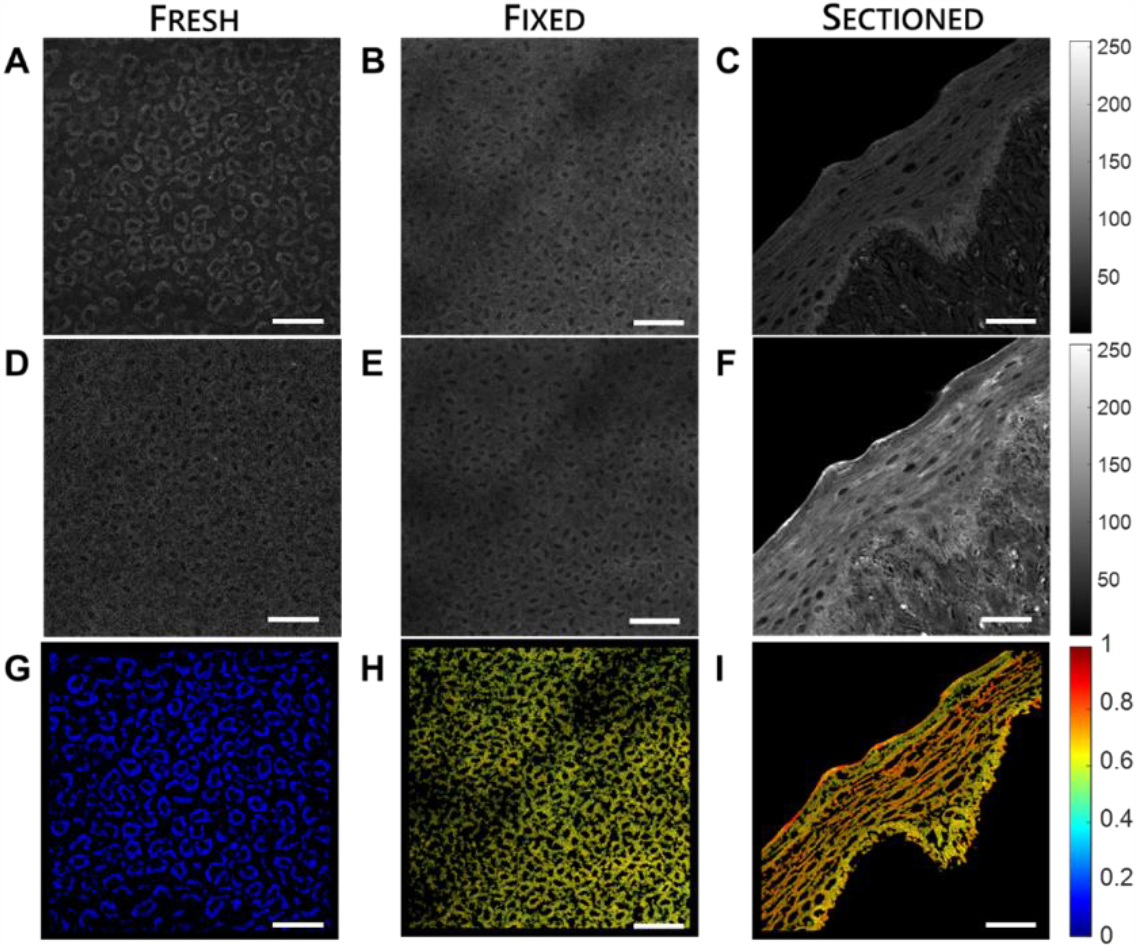
Representative images for bulk fresh (depth of 24 μm), bulk fixed (depth of 40 μm), and sectioned fixed tissue specimens for (A-C) 755 nm excitation, 460/50 nm detection, (D-F) 860 nm excitation, 525/50 nm detection, and (G-I) corresponding optical ratio (I_860/525_/(I_755/460_ + I_860/525_)) coded images. In panel (G) the values correspond to the optical redox ratio (FAD/(NAD(P)H+FAD)). Scale bars are 50 μm. Images on panels C and D have increased contrast for clarity.

The depth dependent mean optical ratio values of each optical section for all ROIs imaged are shown in Fig. 3. The optical (redox) ratio of the fresh tissues is highest at the surface, reaches a minimum at a depth of 20-40 μm, and increases slowly towards the basal layers of the epithelium, resembling a positive parabola or a convex curve. A potential reason for the higher optical redox ratio value at the superficial layers of the freshly excised epithelium is a more oxidized state associated with an increase in FAD as the cells undergo apoptosis [8, 19, 22]. The depth-dependent trend observed in the bulk fixed tissue specimens is not consistent with that observed in the freshly excised specimens. The overall ratio values are significantly higher, exhibiting the lowest ratios at the tissue surface. The depth-dependent increases in these values towards deeper optical sections follow a concave shape. Bulk tissue specimens show a clear continuity in the optical ratio values of adjacent optical sections. This continuity is lost and there is no consistent depth-dependent trend among ROIs after sectioning of the tissue (figure 3C).

**Fig. 3.**
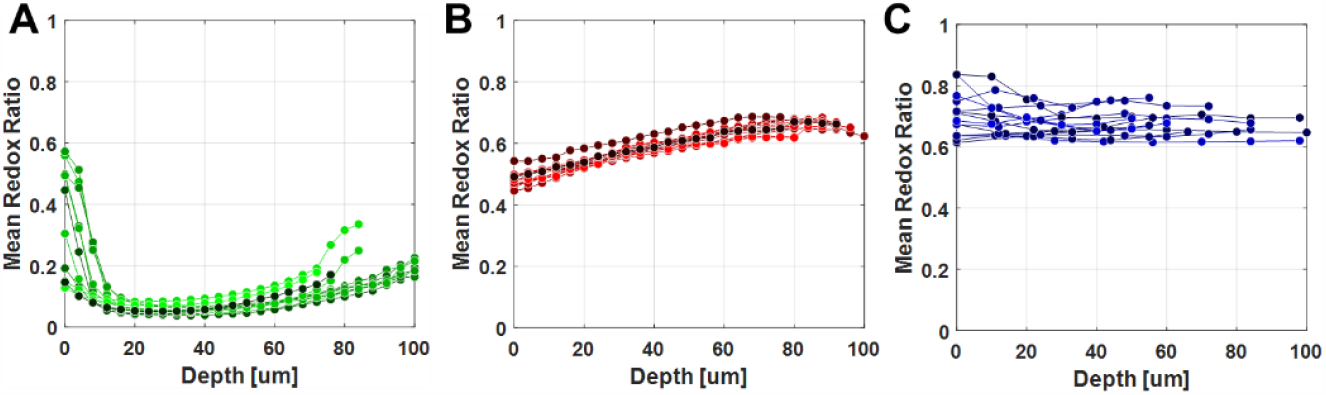
Depth dependent mean optical ratio for each optical section. Each image stack is exemplified with a specific hue of green, red, or blue for (A) fresh, (B) fixed, and (C) sectioned, respectively.

### 3.2 Fixed specimen autofluorescence images lack the cell heterogeneity of fresh tissue

Representative I_755/460_ images showing the intensity variations of bulk fresh and fixed tissue at a depth of 24 μm and 40 μm, respectively, are shown in Fig. 4(A-B). These representative images were selected based on approximate nuclei size. The depth difference is due to hardening of the tissue after formalin fixation and uneven contact with the coverslip. Insets in these two panels are three adjacent cells magnified to highlight their corresponding features. A qualitative assessment of the bulk fresh tissue shows a dark nucleus with lower intensity values, clear intensity variations within the cytoplasmic regions, and darker background regions that permit identification and delineation of each individual cell. The bulk fixed tissue images show a noisier nuclear region, a seemingly homogeneous cytoplasmic region, and loss of distinction between cell borders and interstitial space. The intensity profile of a single-pixel line across two cells in the insets is plotted in figure 4(C-D). The range of values is larger for the fresh tissue cells and there is clear identification of the darker nuclear regions.

**Fig. 4.**
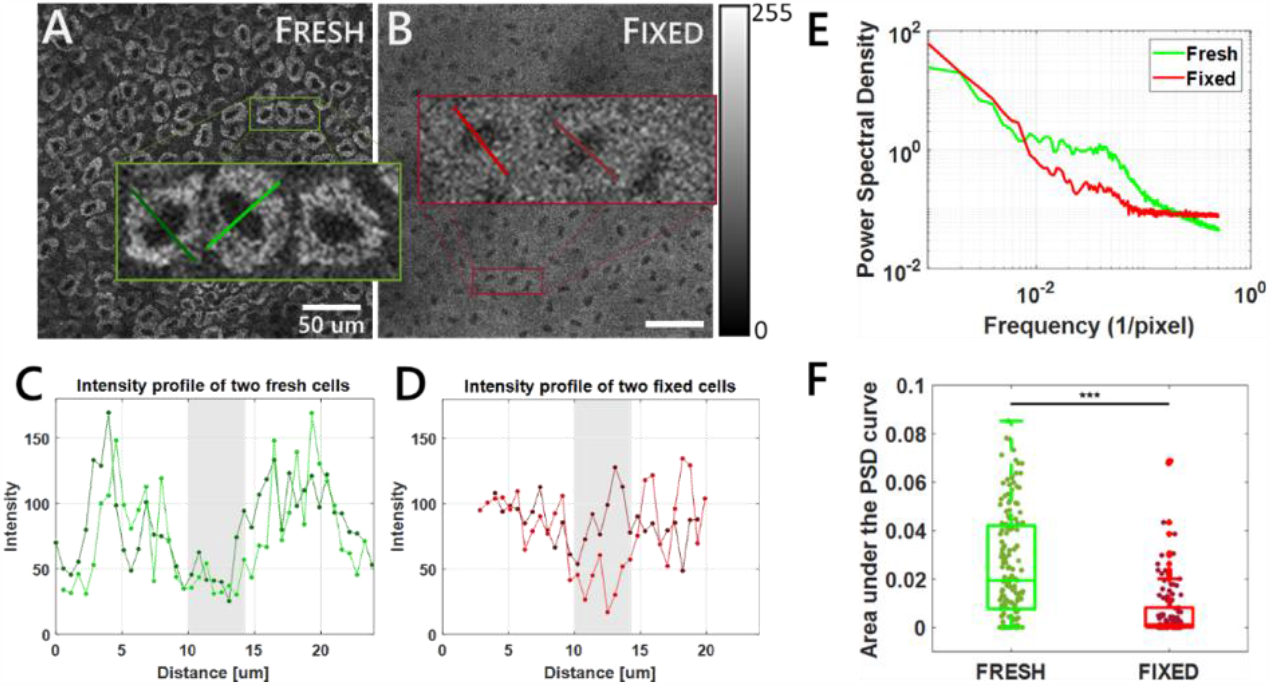
TPEF images of NAD(P)H of a single cell layer of (A) fresh and (B) fixed epithelium at a depth of 24 and 40 μm deep, respectively. Insets are a selection of three adjacent cells within each image. (C-D) Intensity profiles of two representative cells with the nuclei situated at the center of the x axis (shaded region). (C) PSD curves for the images in panels (A) and (B). (F) Area under the PSD curve for frequencies between 7 x 10^−3^ to 2 x 10^−1^ 1/pixel, corresponding to relevant cell features based on sizes of 2-80 μm.

Figure 4E shows PSD curves for representative optical sections of bulk fresh and bulk fixed tissue. The fresh tissue PSD highlights the presence of a significantly higher prevalence of intensity fluctuations with characteristic spatial frequencies between 10^−2^ and 10^−1^ 1/pixel which correspond to sizes of ∼2 to 80 μm. This is quantified by calculating the area under the PSD curve for all optical sections containing nearly full fields of cells in this frequency region (Figure 4F). There are statistically significant differences between both values and, in many cases, the area under the curve for the fixed tissue is very close to zero, as there are few to no contributions of intensity variations, suggesting that the fixation process eliminated intracellular heterogeneity.

### 3.3 Fixation causes significant spectral distortions

Spectral images acquired from fresh, fixed, and sectioned tissues support the presence of significant contributions from fluorophores other than NAD(P)H and FAD. The bulk fresh tissue spectrum at 755 nm excitation is consistent with the emission spectrum of NAD(P)H [44]. The bulk fixed tissue spectrum is broadened significantly, with additional distortions introduced by paraffin embedding and sectioning. It is not clear what portion, if any, of this broadened spectrum can be attributed to NAD(P)H. At 860 nm excitation, spectra from fresh and fixed bulk tissue specimens are consistent with FAD spectra reported in literature [45, 46]. However, distortions can be observed for both excitation wavelengths for the sectioned specimen spectra corresponding to the superficial layers of the epithelium with a peak at ∼540 nm. Even for the bulk fixed specimen, there is a possibility that the spectral overlap with the fresh tissue is coincidental. Peak intensity normalized spectra for both excitation wavelengths are shown in Fig 5.

**Fig. 5.**
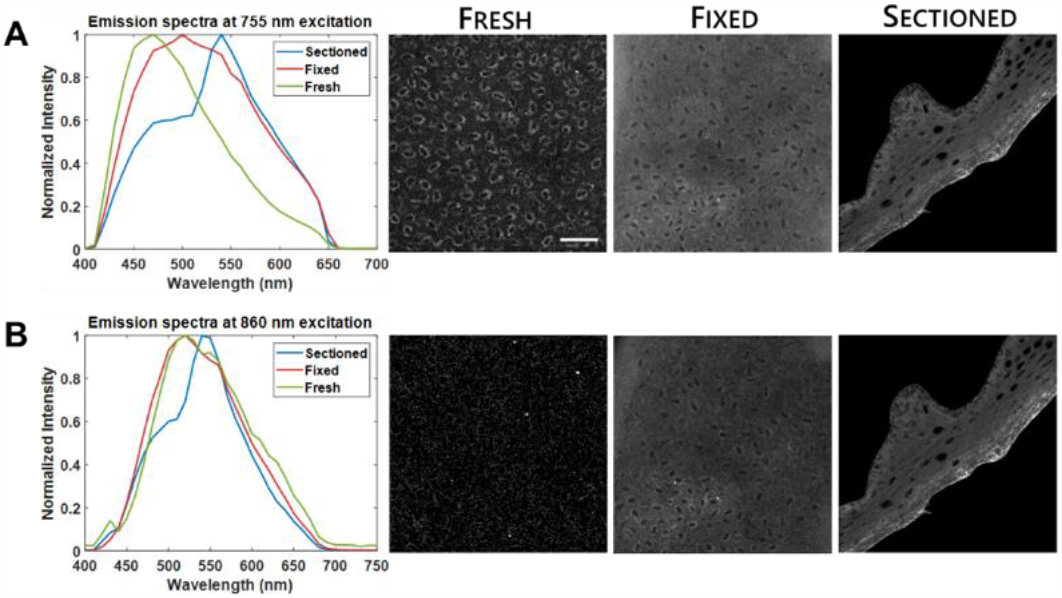
Peak intensity normalized spectra for epithelial cells in bulk fresh, bulk fixed, and sectioned fixed tissue at (A) 755 and (B) 860 nm excitation with the corresponding integrated intensity images. Images have increased contrast for clarity. Scale bar is 50 μm.

**Fig. 6.**
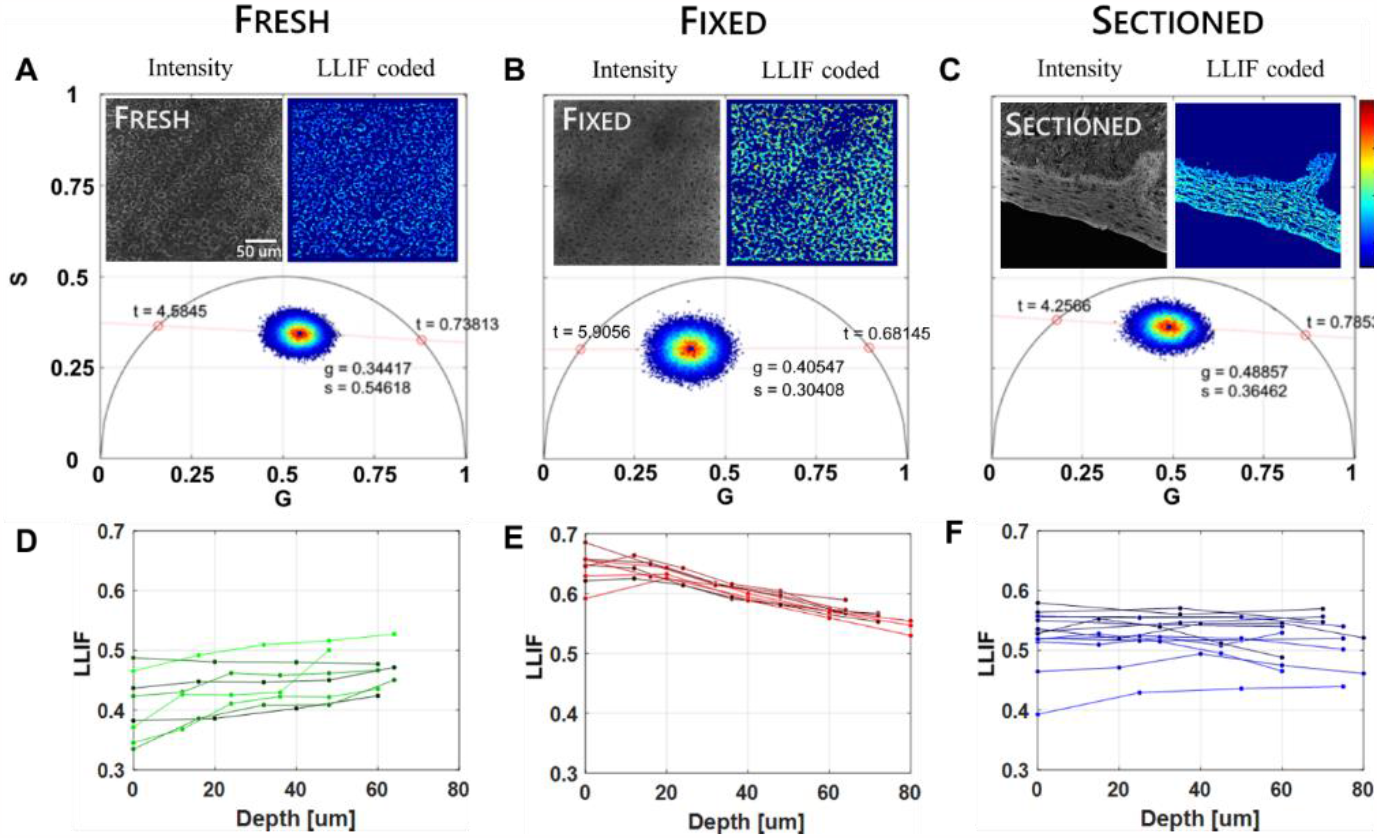
(A-C) Representative NAD(P)H and LLIF coded images of bulk fresh (depth of 36 μm), bulk fixed (depth of 32 μm), and sectioned fixed specimens, respectively, with their corresponding phasor plot. The values for long and short lifetime are the intersects of the fitted line and the universal circle. (D-F) LLIF of bulk fresh, bulk fixed, and sectioned fixed specimens through depth.

**Fig. 7.**
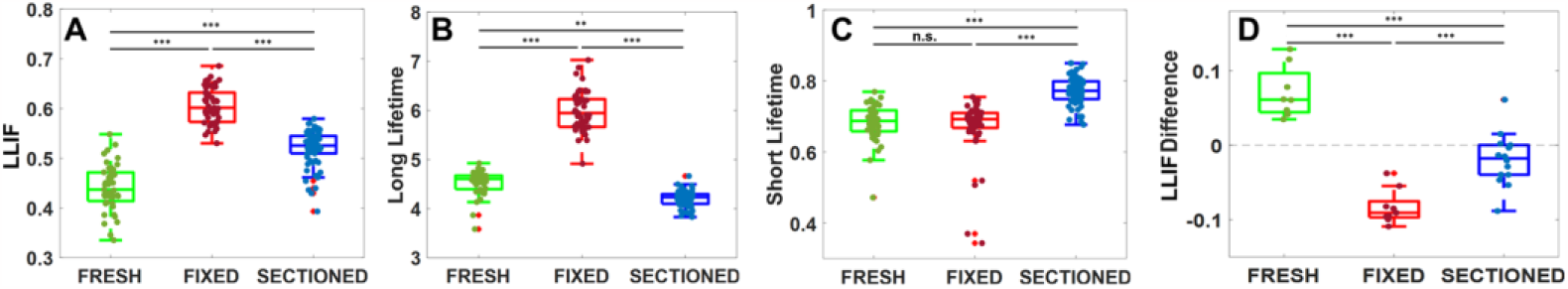
(A-C) Box plots for LLIF, long lifetime, and short lifetime of all optical sections for bulk fresh, bulk fixed, and sectioned fixed specimens, respectively. (D) Difference between the deepest optical section and the most superficial optical section plotted into the different tissue conditions. ** p < 0.001, *** p < 0.0001, n.s. no significance.

**Fig. 8.**
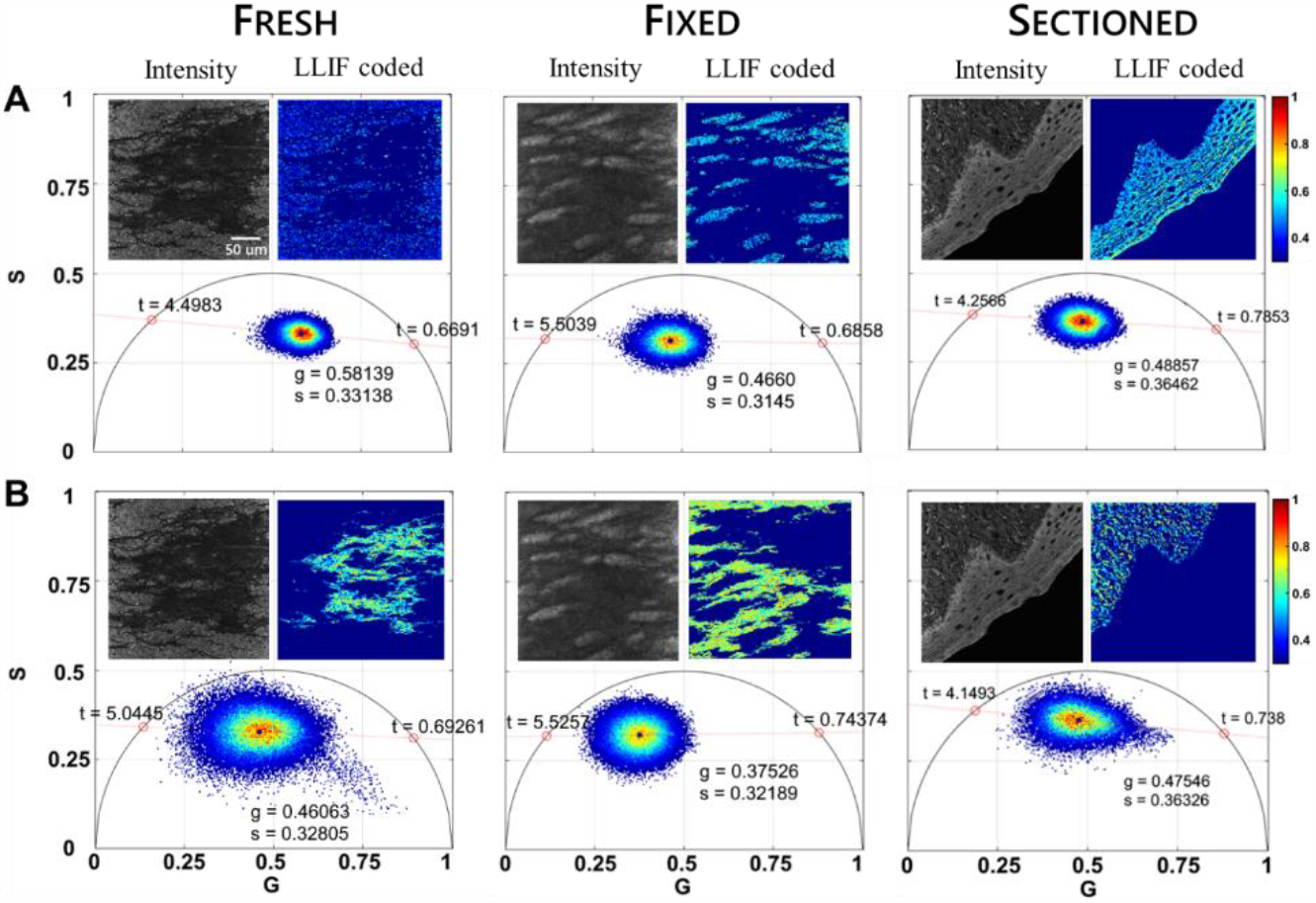
Phasor plots with corresponding intensity and masked LLIF coded images for fresh (depth of 64 μm), fixed (depth of 80 μm), and sectioned tissue. LLIF coded images and phasor plots of (A) cellular regions, and (B) collagen-rich regions.

### 3.4 Fixation and sectioning alter the lifetime characteristics of the fluorescence detected at excitation/emission settings used typically for NAD(P)H imaging

Representative I_755/460_ and LLIF coded images are shown in Fig. 6(A-C) with their corresponding phasor plot for each of the tissue conditions. The optical sections shown for the bulk fresh and fixed images are at a depth of 36 and 32 μm, respectively. The two *t* values in the phasor plots are the estimated long and short lifetime of the fluorescent molecules within the corresponding optical sections. Figure 6(D-F) shows the depth-dependence of the long lifetime intensity fraction for all ROIs imaged. For the freshly excised specimens, these are the NAD(P)H bound fraction values, which increase as a function of depth, yielding a positive slope as the general trend. The opposite is seen in the bulk fixed specimens, where a negative slope is observed in the depth dependence of the LLIF. This parameter exhibits only small and inconsistent depth-dependent variations in sectioned fixed samples.

There are statistically significant differences for LLIF for all conditions (Fig. 7A). Examination of the long lifetime contributions for freshly excised biopsies are consistent with the typical values of bound NAD(P)H found in literature [14] and range from 3.59 to 4.93 ns. There is a significant increase in the long lifetime when analyzing bulk fixed specimens with values ranging from 4.91 to 7.03 ns, while the range of values for the long lifetime of sectioned specimens is 3.84 to 4.37 (Fig. 7B). On the other hand, there is no significant difference in the short lifetimes for bulk fresh and bulk fixed specimens (Figure 7C). However, the short lifetime value for sectioned specimens is statistically higher from the that of the bulk specimens, with a range from 0.69 to 0.81 ns. In figure 7D we quantify the difference between the long lifetime intensity fraction of the deepest and most superficial optical sections to highlight the statistically significant differences in the depth-dependence of this parameter among all tissue conditions. The change in the trends for all optical metabolic function metrics suggests that the fixation and tissue processing for paraffin embedding and sectioning poses significant limitations in the ability to assess metabolic function from endogenous TPEF intensity and lifetime images. This is further supported when observing signals from the basal layers of the epithelium that are rich in collagen.

Figure 8 shows phasor plots with their corresponding intensity and LLIF images from tissue depths of 64 and 80 μm for fresh and fixed tissue, respectively. Lighter regions in the intensity images correspond to NAD(P)H signal for fresh specimens and darker regions show weak fluorescence that is emanating from collagen. To highlight the differences between the phasors and the centroid location from cellular and collagen regions, a mask of collagen positive pixels (SHG) was created for the same optical section. Figure 8A shows the traditional masking, which analyzes I_755/460_ signal and removes background as described in the methodology. Figure 8B shows the SHG positive, collagen-rich regions. The phasor of the collagen rich pixels is noisier than that of the cell pixels because the signal is weaker; however it is clear that the centroid is displaced relative to that of the cells and the lifetimes tend to be higher. The cell phasor of the fixed bulk specimen shows shifts consistent with the changes reported in Fig. 5-6 and appears much more similar to the phasor of the collagen positive pixels. The overlap is even more significant in the sectioned tissues, with both the short and long lifetimes of the cellular and collagen regions appearing very similar. These shifts highlight the presence of signals in the sectioned tissues that are not necessarily of the same origin as the fresh tissues. Unfortunately, assessing what contributions can be attributed to endogenous vs crosslink fluorescence is not possible from these studies.

## 4. Discussion

The main goal of this study was to determine whether metabolic interpretations from fresh tissue TPEF images acquired with excitation/emission settings attributed to NAD(P)H and FAD are maintained post formaldehyde-based fixation and histology processing. Evaluation of the preservation of optical metabolic function readouts was performed through imaging of the same tissues in three different conditions: freshly excised, formalin fixed, and sectioned after paraffin embedding. The TPEF-related metrics used for comparison were the optical (redox) ratio (I_860/525_/(I_755/460_ + I_860/525_)), intra and intercellular intensity variations, fluorescence lifetime, and spectral data. Our results indicate that the fixed tissue images cannot be assumed to reflect reliably the metabolic function of the corresponding fresh tissues prior to fixation.

The TPEF intensity variations attributed to metabolic heterogeneity and the interstitial space delineating individual cells are lost after fixation in our studies (Fig. 4). The higher intensities in the 525/50 nm detection band for fixed tissue cause a shift towards higher optical ratio values and the different optical sections of the bulk fixed tissue show an opposite trend through depth when compared to fresh tissue (Fig. 3). This trend is lost in the sectioned tissue analysis (Fig 3C). Thus, the optical ratio images of the fixed specimens in our study do not convey the same differences in metabolic function as the corresponding optical redox ratio images of fresh tissues.

Spectra acquired at 755 nm excitation highlight a significant broadening upon fixation and sectioning relative to the spectrum attributed to NAD(P)H TPEF from the freshly excised epithelium (Fig. 5). NAD(P)H could contribute to the spectra acquired from fixed and sectioned tissues, but its intensity relative to additional autofluorescence likely arising from the cross-links that are formed is not possible to assess from these measurements. While fixation increases the fluorescence detected at 860 nm excitation, the spectrum of the emitted fluorescence is not modified relative to that acquired from the freshly excised tissues. However, we do detect some differences upon paraffin embedding and sectioning. These investigations suggest that the signals gathered at both excitation/emission wavelength ranges we explored are likely influenced significantly by the cross-links generated during different tissue processing steps.

This conclusion is supported further by the lifetime measurements. The depth-dependent variations of lifetime measurements are not preserved reliably after fixation and sectioning (Fig. 6). The significant phasor differences between cellular and collagen regions in the fresh tissues at 755 nm excitation appear minimal in the sectioned tissues, suggesting similar dominant contributions from these two regions in FFPE samples. The main fluorophore within the collagen rich region cannot be NAD(P)H, further supporting a prominent role for the signal emanating from newly formed crosslinks in both stromal and epithelial areas.

Fixation has been shown to impact the tissue spectral emission in a manner that is dependent on the type of tissue and the time of fixation. While in some cases major differences are reported within minutes [33] or 24 hours [47] from the onset of fixation, in other studies significant shifts are observed throughout four days of fixation [48]. In the latter study, which focused on 800 nm TPEF of mouse skeletal muscle tissues, spectra with distinct peaks at 470nm and 550 nm from fresh tissues, were replaced by broader spectra, which in principle could represent a different balance in contributions from NAD(P)H and FAD or additional fluorescence from formed cross-links [48]. The overall detected fluorescence intensity increased significantly during the last two days of fixation. In a different study, single photon excited fluorescence spectra at 488 nm from FFPE mouse colon tissues appeared consistent with FAD emission, but attribution of the signal to FAD was not validated any further [49]. At 337 nm excitation, significant distortions were observed following formalin fixation of breast tissues [50]. These changes were attributed to quenching of the fluorescence with a 390 nm emission peak attributed to collagen and elastin by the crosslinks formed during fixation. It was presumed that the spectrum with a single peak at 455 nm following fixation was associated with NAD(P)H, with the intensity enhanced preferentially in cancer tissues because they were expected to be impacted much more prominently by dehydration compared to the lipid rich normal tissues. As these were made with a point probe, only bulk tissue considerations could be made. While the impact of hemoglobin leaching from the tissue during the multi-day fixation processing was identified as a potential source of intensity changes, its impact on spectral changes was not considered. The option of having cross-links contribute to the 455nm peak was also not discussed. These studies highlight that fixation does impact the spectral characteristics of the emitted fluorescence, but the extent to which these fluorescence changes originate from simple changes in the signals that were present pre-fixation or whether new species are potential significant contributors was not assessed robustly.

TPEF images acquired from mouse muscle samples at excitation/emission wavelengths similar to those used in our study highlight differences, even after 5 min of fixation with 4% paraformaldehyde (PFA) [31]. These changes were attributed to a consistent increase in the autofluorescence intensity detected at 860 nm excitation, 515-535 nm emission (where FAD TPEF was detected in freshly excised tissues) and a somewhat inconsistent decrease in the autofluorescence detected at 750 nm excitation, 435-485 nm emission (where NAD(P)H TPEF was detected from freshly excised tissues). These trends are consistent with our observations and in fact highlight the possibility that there are other significant contributors to the fluorescence signal detected especially under excitation/emission wavelengths consistent with flavin fluorescence. Interestingly, despite this observation the investigators continued to assess the optical ratio in FFPE slides of muscle sections from mice of different ages assuming that NAD(P)H and FAD are the dominant signals. We note that fixation time of such samples was approximately 16 hours.

It has been reported that there is an increase in mean lifetime values of NAD(P)H due to the fixation process [33]. This increase was noticed in our experiments as well. However, this is attributed primarily to an increase in the long lifetime and LLIF values of bulk fixed specimens, and an increase in the short lifetime of fixed sectioned specimens. Thus, it is not clear that the signal detected at I_755/460_ from fixed specimens is indeed representative of NAD(P)H. The significant overlap of the lifetime phasor signatures corresponding to the stromal and the epithelial regions of the FFPE sections further supports the possibility that the dominant detected contributions are not attributed to NAD(P)H

Interestingly, Chacko *et al*. [33] reported preservation of the subcellular localization of NAD(P)H-associated lifetime contrast before and after 24 hour fixation of cell cultures, even though there was a significant increase in the mean lifetime of fixed cells. The impact of fixation also was different for healthy and cancerous cells or cells treated with different metabolic inhibitors. However, differences among different cell populations were preserved. Interestingly, when the impact of fixation was considered on fluorescence intensity detected at wavelengths corresponding to FAD (740 nm or 890 nm excitation, 525/50 emission) fluorescence, it was found that fixation did not allow preservation of differences identified in corresponding live cell culture measurements. In fact, the trends detected in the FAD-associated settings were similar to the NAD(P)H-associated signal. This was attributed to a spectral shift in the NAD(P)H signal following fixation. Potential contributions to the detected signals from fixation-induced crosslinks were not considered.

Normal and cancer mammary tissues were also considered in this thorough study. The preservation of subcellular contrast was not reported after fixation for these tissues. Thus, it is not clear if fixation may impact fluorescence localization differently in cells and tissues. However, differences in the mean lifetimes of normal and cancer breast tissues were preserved following fixation. The excitation/emission wavelengths at which the reported lifetimes were recorded were not indicated explicitly in the manuscript. We also note that that these results were most likely from unsectioned fixed tissues. Further, it appears that mostly adipose tissue contributed to the normal tissue signal, while a wider mixture of collagen, cancer cells and adipocytes contributed to the reported cancer tissue signals. To further assess the impact of paraffin embedding and sectioning, the authors presented intensity and lifetime images acquired at 740 nm excitation but over a 100 nm emission bandwidth centered at 460 nm (attributed to NAD(P)H), along with second harmonic generation images from pancreatic tissue microarrays. Interesting differences were identified in the mean lifetimes between cancer and normal pancreatic tissues. These tissue sections were very rich in collagen, as highlighted by the SHG images, and contributions from the collagen cross-links to the signal attributed to NAD(P)H fluorescence were not considered. Thus, while the presence of lifetime contrast between normal and cancer specimens upon fixation is supported by the results, the origin of the signal that leads to this differentiation may not necessarily be NAD(P)H.

The lack of consistency in some of the reported results further supports that the fluorescence intensity and lifetime signatures from fixed specimens cannot be reliably attributed to report on metabolic function. For example, while MCF10A breast epithelial cells were found to exhibit a significantly higher NAD(P)H-associated mean lifetime by Chacko et al. [33] following fixation, no lifetime differences were found for the same cells before and after fixation by Conklin et al. [32] at wavelengths associated with images acquired at either 780 nm or 890 nm excitation. The cells were embedded in a collagen gel in the latter study. In addition, even though the same breast tumor model was used in these two studies (PyVmT), a longer lifetime was reported for the tumor cells by Conklin et al, while a shorter mean lifetime was reported by Chacko et al. The former study focused its comparisons to ROIs containing only epithelial cells, while this was not the case in the study by Chacko et al. The results were also likely impacted by the use of FFPE slides (Conklin et al) vs fixed tissues (Chacko et al). Spectra were also reported at these two excitation wavelengths from an FFPE tissue section acquired from the same breast cancer model by Conklin et al. At both of these wavelengths, the spectra appear similar to what we report in our study (Fig.5) with the major peak exhibiting a maximum at 530-540 nm.

Following fixation, there appears to be a consistent increase in the signal collected at wavelengths where flavin autofluorescence would be typically collected among all studies. Fixation also leads to an increase in the fluorescence lifetime of signals detected at wavelengths where NAD(P)H signals are typically excited. In addition, several studies highlight that potentially diagnostically useful information is preserved in TPEF intensity and lifetime images of fixed tissues and FFPE tissue sections. However, our studies indicate that great caution should be exerted when attributing the signals detected to NAD(P)H and FAD. The spectral data acquisition for I_755_ showed broadening of the spectrum and a shift in peak emission in fixed specimens compared to the fresh samples, which suggest the presence of additional fluorophores with emission bands overlapping NAD(P)H emission. Our bulk fixed tissue analysis revealed elimination of the interstitial space between cells in the resulting images for both I_755/460_ and I_860/525_. The significant overlap in the lifetime signatures of collagen rich and epithelial cell rich regions of FFPE tissue sections also support that fixation products dominate the detected endogenous fluorescence signals. However, we note that in our studies tissues were fixed with 10% formalin for a week. The concentration of the fixative as well as the duration of fixation will likely impact outcomes. However, other fixation protocols, such as ones based on milder sucrose fixation and cryosectioning may be better suited for preserving endogenous fluorescence signals that are more directly linked to NAD(P)H and FAD.

## Acknowledgments

The authors wish to thank Dr. Michael Esmail for assistance with procuring the animal tissues used in these studies.

## Data availability

The data presented in this paper can be made available from the corresponding author upon reasonable request.

